# Learning Peptide Recognition Rules for a Low-Specificity Protein

**DOI:** 10.1101/2020.06.02.131086

**Authors:** Lucas C. Wheeler, Arden Perkins, Caitlyn E. Wong, Michael J. Harms

## Abstract

Many proteins interact with short linear regions of target proteins. For some proteins, however, it is difficult to identify a well-defined sequence motif that defines its target peptides. To overcome this difficulty, we used supervised machine learning to train a model that treats each peptide as a collection of easily-calculated biochemical features rather than as an amino acid sequence. As a test case, we dissected the peptide-recognition rules for human S100A5 (hA5), a low-specificity calcium binding protein. We trained a Random Forest model against a recently released, high-throughput phage display dataset collected for hA5. The model identifies hydrophobicity and shape complementarity, rather than polar contacts, as the primary determinants of peptide binding specificity in hA5. We tested this hypothesis by solving a crystal structure of hA5 and through computational docking studies of diverse peptides onto hA5. These structural studies revealed that peptides exhibit multiple binding modes at the hA5 peptide interface—all of which have few polar contacts with hA5. Finally, we used our trained model to predict new, plausible binding targets in the human proteome. This revealed a fragment of the protein *α*-1-syntrophin binds to hA5. Our work helps better understand the biochemistry and biology of hA5, as well as demonstrating how high-throughput experiments coupled with machine learning of biochemical features can reveal the determinants of binding specificity in low-specificity proteins.

## Introduction

Up to 40% of protein-protein interactions are mediated by globular domains that recognize a short, linear fragment of their interaction partner. ^1,2^ Such protein-peptide interactions play key roles in processes ranging from from signaling networks to biological phase transitions.^2,3^ Understanding such systems therefore requires knowing which proteins recognize which peptides under what conditions.^2,4^

Protein-peptide interaction interfaces exhibit a wide range of specificity. For some proteins, one can describe specificity using a simple binding motif that encodes the amino acid(s) recognized at each site in the peptide.^5,6^ One can predict protein targets by searching for matching sequences within the proteome. ^6^ Some proteins deviate from this highly specific paradigm, requiring more sophisticated approaches. For example, many PDZ binding domains exhibit binding “multi-specificity”, in which peptide preference must be represented as a handful binding motifs.^7,8^ Predicting interaction targets for such proteins is more difficult than for proteins with single binding motifs, but the same basic logic applies: search the proteome for sequences that match the binding motifs.

Even more extreme cases exist, such as S100 proteins. Members of this family of calcium-activated signaling proteins play important roles in a wide range of critical cellular processes such such as innate immunity, cell-cycle regulation, and inflammatory signaling.^9,10^ S100s bind to short, linear peptide regions of target proteins, modulating their activity (Fig 1).^9–17^ Defining the peptide recognition rules of S100 proteins has, however, proven extremely difficult, as target peptides lack sufficient similarity to be usefully represented as a binding motif.^18,19^ That said, the low-specificity of the S100 protein family does not equate to *no specificity*. Specific targets within the highly-variable sets of S100 binding partners appear to be evolutionary conserved over hundreds of millions of years, even as the interface acquired mutations.^18^

**Fig 1.**
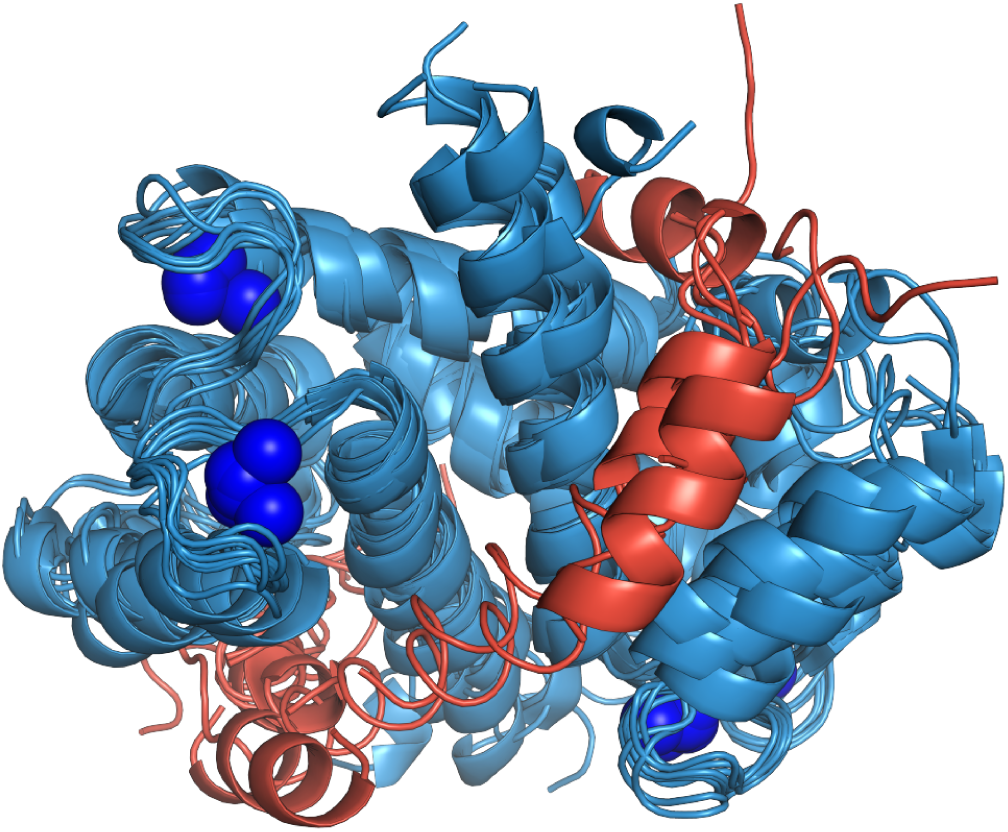
S100 proteins interact with peptides in a canonical peptide binding interface in a calcium-dependent manner. Structure shows an alignment eight different S100 structures (PDB IDs: 3IQQ, 1QLS, 3RM1, 2KRF, 4ETO, 2KBM, 1MWN, 3ZWH) with peptides bound at the canonical interface (red). Calcium ions are shown as blue spheres.

Here we endeavor to dissect the specificity of a representative S100 protein: human S100A5 (hA5). This protein likely plays a signaling role in olfaction. ^20,21^ It interacts with a diverse set of peptide targets with no obvious sequence motif.^17–19^ Instead of representing hA5’s peptide specificity with a binding motif, we represented it using readily calculated biochemical properties of each peptide. We used machine-learning to train a model against a recently released quantitative phage display dataset ^22^, finding that we were able to reproduce the phage display data and several measured peptide binding interactions formed by hA5. We then used this model both to understand what biochemical features hA5 recognizes and to predict new binding targets. Our results demonstrate that it is possible to gain insights into the rules that define binding specificity, even for proteins with extremely low specificity. The software we developed for this purpose—HOPS: <u>H</u>unches from <u>O</u>regon about <u>P</u>eptide <u>S</u>pecificity—is available for download (https://github.com/harmslab/hops).

## Results

### Peptide sequence is insufficient to describe specificity

Previously, we collected a high-throughput, phage-display dataset for hA5 interacting with ≈ 40, 000 random 12-mer peptides.^22^ We did two parallel panning experiments: one in the absence of competitor (Fig 2A), the other in the presence of a competitor peptide that binds at the site of interest (Fig 2B). We then used high-throughput sequencing to quantify the frequencies of peptides in both the “conventional” and “competitor” samples (*f*_*conventional*_ and *f*_*competitor*_, respectively). Peptides that bind at the site of interest are depleted preferentially in the competitor sample. This can be quantified by *E* = *ln*(*f*_*competitor*_*/f*_*conventional*_), meaning that *E* < 0 corresponds to a peptide that binds at the site of interest. We found we that peptides with *E* < −1.37, corresponding to a four-fold decrease in frequency with the addition of competitor, could be distinguished from zero with a false-discovery rate of 0.05. Throughout our analysis, we therefore use a cutoff of *E* = −1.37 for peptides that bind. We first sought to determine if sequence-level rules were sufficient to describe the specificity of peptides enriched by hA5 in the phage display enrichment experiment. We took the 3, 574 unique peptide sequences with *E* < −1.37 calculated a position-specific weight-matrix.^23^ This approach revealed extreme variability across positions in the peptide (Fig 2C).

**Fig 2.**
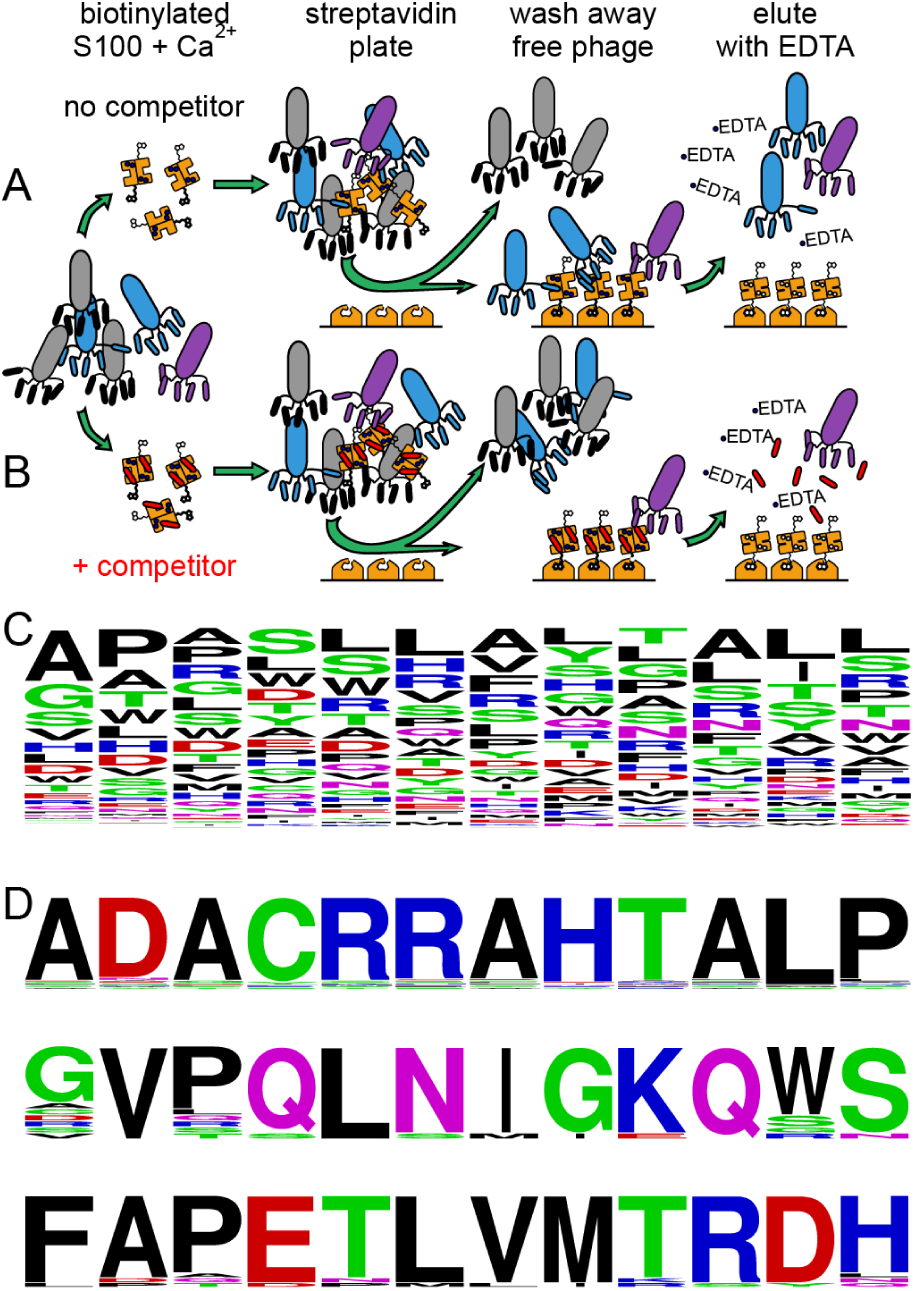
Interacting peptides can be identified using phage display. Panels A and B) Rows show two different experiments, done in parallel, for each protein. Biotinylated, *Ca*^2+^-loaded, hA5 is added to a population of phage either alone (row A) or in the presence of saturating competitor peptide (row B). Phage that bind to the protein (blue or purple) are pulled down using a streptavidin plate. Bound phage are then eluted using EDTA, which disrupts the peptide binding interface. In the absence of competitor (row A), phage bind adventitiously (purple) as well as at the interface of interest (blue). In the presence of competitor (row B), only adventitious binders are present. C) Sequence logo for all peptides in the phage display dataset for which *E* < −1.37. Each position is highly variable in the position-weight-matrix. D) Frequency sequence logos representing three of the 28 peptide clusters identified using DBSCAN.

We next used a clustering approach to identify a set of motifs representing a profile of “multi-specificity” for hA5. Similar approaches have worked for other multi-specific proteins, identifying a small set of motifs sufficient to describe binding preferences. ^8,24^ We clustered enriched peptides by Damerau-Levenstein distance using the DBSCAN algorithm.^25–27^ We then generated position-specific-weight-matrices for each cluster. Only 0.4% of peptides were placed in clusters; the remainder were placed into singleton clusters. The resulting clusters were highly diverse. Fig 2D shows three of the identified clusters: there is little sequence commonality between the three clusters. This extreme “multi-specificity” of hA5 extends beyond a small set of motifs easily represented by position-weight matrices. Thus, we concluded that simple sequence-based rules were insufficient to identify the key determinants of specificity in hA5 or to construct a predictive model for binding partners.

### Supervised machine learning can be used to train a predictive model of enrichment

We sought an alternate approach to simple sequence-based metrics. Inspired by the literature on characteristic biochemical properties of intrinsically-disordered proteins, ^28^ we hypothesized that the biochemical features of peptide targets could be used to construct a predictive model of of peptide enrichment. For each peptide sequence identified in the phage display dataset, we calculated a set of 57 features covering an array of biochemical properties. We included properties such as hydrophobicity, Chou-Fasman *α*, accessible surface area, isoelectric point, and net charge (a full list of all predicted features is available in Table S1). We calculated each feature in sliding windows across the peptide sequence, resulting in a final set of 4,446 features for each individual peptide (Fig. 2A).

We then trained a Random Forest model to reproduce our phage display *E* values using these features as inputs.^29^ Prior to training the model, we withheld 10% of the data to use as a test set. We then optimized the nuisance parameters in our model—which features to include, whether to apply a sliding window, and the number of estimators—using k-fold cross-validation. The best model we identified used sliding windows, included the full set of 57 base features, and used 30 estimators (Fig 2B). We then trained our model against the complete training set and measured its predictive power using the previously withheld 10% test set. This yielded a final 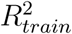 of 0.973. Overall, the model reproduced the test set phage display *E* values well, giving 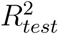 of 0.867 (Fig. 2C). There was a systematic deviation between the predicted and measured values of *E* for the highest and lowest values: the slope between *E*_*predicted*_ and *E*_*measured*_ was also 1.16 rather than 1.00 (Fig 2C). This suggests that there are features important for the highest and lowest *E* values not fully captured by our model.

We also tested the utility of using the model to predict whether *E* for a peptide was expected to be above or below the *E* cutoff of −1.37. We calculated a Receiver Operator Characteristic (ROC) curve for the classifier, plotting the true positive rate against the false positive rate. This yielded a highly predictive model, with an area under the curve of 98.9. This give us high confidence in using this model to classify enriching peptides.

We validated our model by reproducing the known binding of four S100 peptide targets that we had previously studied using isothermal titration calorimetry (ITC).^18^ Of these, three bind to hA5 and one does not. We used the model to predict whether the known peptides would bind to hA5. For peptides longer than 12 amino acids, we calculated the score for all possible contiguous 12-mers and took the best score as our predicted *E* value. The model predicted binding of the peptides NCX1, A5cons, and A6cons, while predicting that the SIP peptide would not bind (Table 1). These predictions accurately recapitulate the known pattern of binding for all four peptides.

**Table 1:**
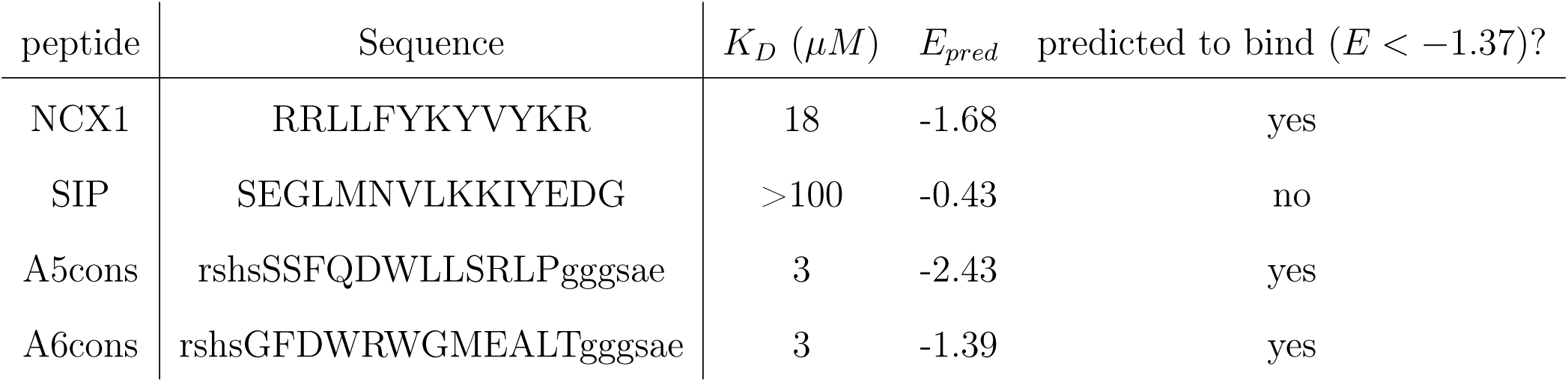
Dissociation constants and model predictions for known peptide targets. Data for the known target peptides used in our previous study. ^18^ NCX1 and SIP are fragments of human proteins. A5cons and A6cons were identified as consensus sequences from an earlier phage display experiment. The lowercase flanking sequences “rshs” and “gggsae” come from the M13 phage coat protein. *K*_*D*_ and predicted E value (*E*_*pred*_) are shown. The statistically significant *E* cutoff for hA5 is −1.37.

### Model classifies peptides based on hydrophobicity and shape complementarity

We next asked what aspects of the peptides were recognized by the trained model. We quantified the contribution of each feature and peptide position to the predicted *E* as measured by node impurity (see methods). We found that no one feature or peptide position dominated the prediction (Fig 3E). To summarize the data, we pooled features based on their chemical similarity. For example, we pooled side chain volume and beta-chain knob propensity,^30,31^ along with a variety of other terms, into “geometry”. We pooled predicted charge and number of hydrogen bonds, on the other hand, into “polar” interactions. A full list of the individual features and their bins is given in Table S1.

**Fig 3.**
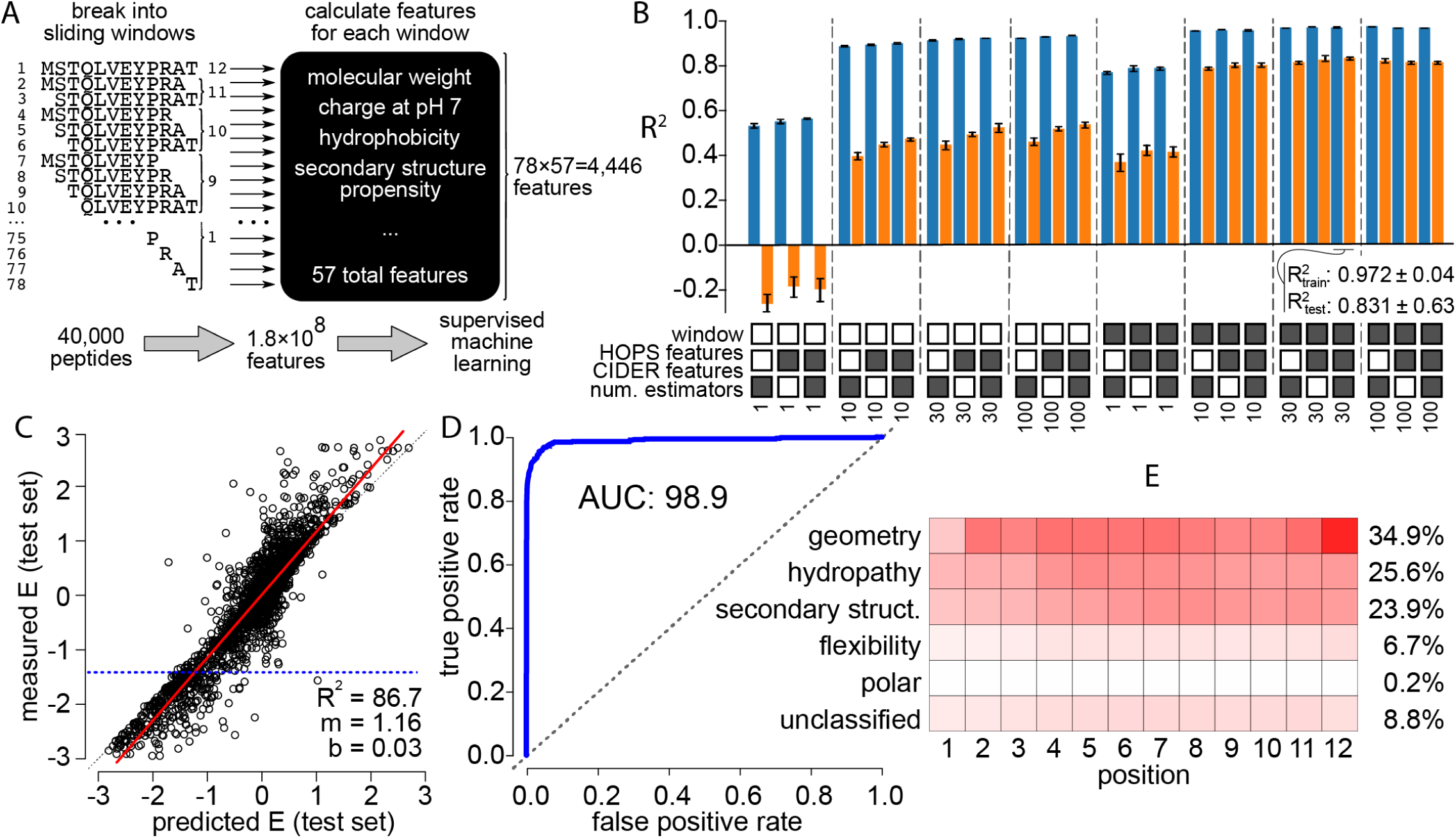
Machine learning model predicts phage display enrichment. A) Diagram of the process for training the machine-learning model. Peptides are broken into sliding windows and a set of predicted biochemical features is calculated for each window. These are the features used in the machine-learning model. B) We found best model input parameters using cross-validation. Pairs of bars represent the average 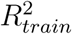 (blue) or 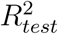 (orange) for 10-fold cross-validation replicates of the data using the model parameters below. Square indicates whether the feature was used in the model (filled) or not (empty). “Window”: whether sliding windows were used. “HOPS” and “CIDER” features are listed in Table S1.“Num. estimators” is number of estimators included in the Random Forest. The 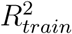 and 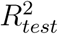 are indicated for the chosen model. C) Points are individual peptides. Red line is the a linear regression between the predicted *E* and measured *E* for each peptide in the test set. Dashed line blue line indicates the threshold below which we can measure enrichment (*E* = −1.37). D) ROC curve for classifying peptides as above or below the *E* cutoff. The area under the curve is shown on the plot. E) Heat map shows the contribution of each site (position 1-12) and aggregated chemical feature (top-to-bottom) to the final model. Color indicates relative contribution from red (strong) to white (no contribution). The marginal contribution of each chemical feature is shown to the right of the plot. Table S1 describes which chemical features went into which aggregate bins.

We plotted the relative contribution of each property as a function of peptide position (Fig. 3E). Each site contributed almost equally to the predicted enrichment (Fig. 3E). Meanwhile, different molecular properties had radically different contribution levels. Geometry, hydropathy, and secondary structure propensity dominated the predictive power of the model. In contrast, polar contacts—often a strong determinant of specificity—had almost no predictive power (Fig. 3E). These results are consistent with binding being determined by shape complementarity and hydrophobic interactions at the interface.

### A high resolution crystal structure reveals binding site interaction variability in hA5

To test the hypothesis that peptide binding was determined by shape complementarity and hydrophobicity, we sought to co-crystallize hA5 with a bound peptide. Despite multiple attempts, however, we were unable to grow hA5 crystals in the presence of peptides, nor to soak peptide targets into crystals grown in the absence of peptide. We did, however, discover a new crystal form of calcium-bound hA5 in the absence of peptide. This crystal diffracted to 1.25 *Å*—the highest resolution crystal structure so far determined for hA5 (PDB: 6WN7, Fig. 4A,B Table 2). The structure contains three homodimers (chains AB, CD, and EF) with similar global structure (0.98-1.78 *Å* RMSD across all *C*_*α*_), but that form different interactions with one another (Fig. 4A yellow ovals). An overlay shows good agreement for individual chains from a previously-solved 2.6 *Å* crystal structure and an NMR structure. The RMSD was 2.0 *Å* over all *C*_*α*_, with the regions of highest variation being Ser43-Glu49 and the C-terminal helix (Fig. 4B). The position of the homodimer partner chains also show variability, with shifts of 1-3 *Å* between structures (Fig. 4C).

**Table 2:** Crystallography data collection and refinement statistics. ^*a*^Resolution cutoff was applied using *CC*_1/2_ > 0.3. ^*b*^Resolution at which < *I/σ* > falls to 2.0. ^*c*^Data may be twinned, inhibiting further model improvement.

|  |  |
| --- | --- |
| ***Data collection*** | PDB code: 6WN7 |
| Structure | Human S100A5 C43S, C79S |
| Space group | $P3_2$ |
| Unit cell a, b, c ($\text{\AA}$), $\alpha, \beta, \gamma$($^\circ$) | 76.3, 76.3, 84.2, 90.0, 90.0, 120.0 |
| Resolution ($\text{\AA}$) | 51.98 – 1.25(1.32–)<sup>a</sup> |
| Completeness (%) | 99.8 (99.8) |
| Unique reflections | 151437 (4561) |
| Multiplicity | 10.6 (6.48.4) |
| $R_{meas}$ (%) | 8.1 (115.8) |
| $CC_{1/2}$ | 1.0 (0.48) |
| $\langle I/\sigma \rangle$ | 16.2 (2.1) |
| $\langle I/\sigma \rangle \sim 2.0(\text{\AA})^b$ | 1.24 |
| ***Refinement*** |  |
| Resolution range ($\text{\AA}$) | 26.0-1.25 |
| R-factor (%) | 26.2<sup>c</sup> |
| R-free (%) | 29.4<sup>c</sup> |
| Protein residues | 159 |
| $Ca^{2+}$ | 12 |
| Water molecules | 498 |
| RMSD lengths ($\text{\AA}$) | 0.009 |
| RMSD angles ($^\circ$) | 1.1 |
| *Ramachandran plot*<sup>c</sup> |  |
| $\varphi, \psi$ – *Preferred* (%) | 98.66 |
| $\varphi, \psi$ – *Allowed* (%) | 1.34 |
| $\varphi, \psi$ – *Outliers* (%) | 0.0 |
| *B-factors* |  |
| $\langle \text{Protein atoms} \rangle$ ($\text{\AA}^2$) | 24 |
| $\langle \text{Waters} \rangle$ ($\text{\AA}^2$) | 33 |

**Fig 4.**
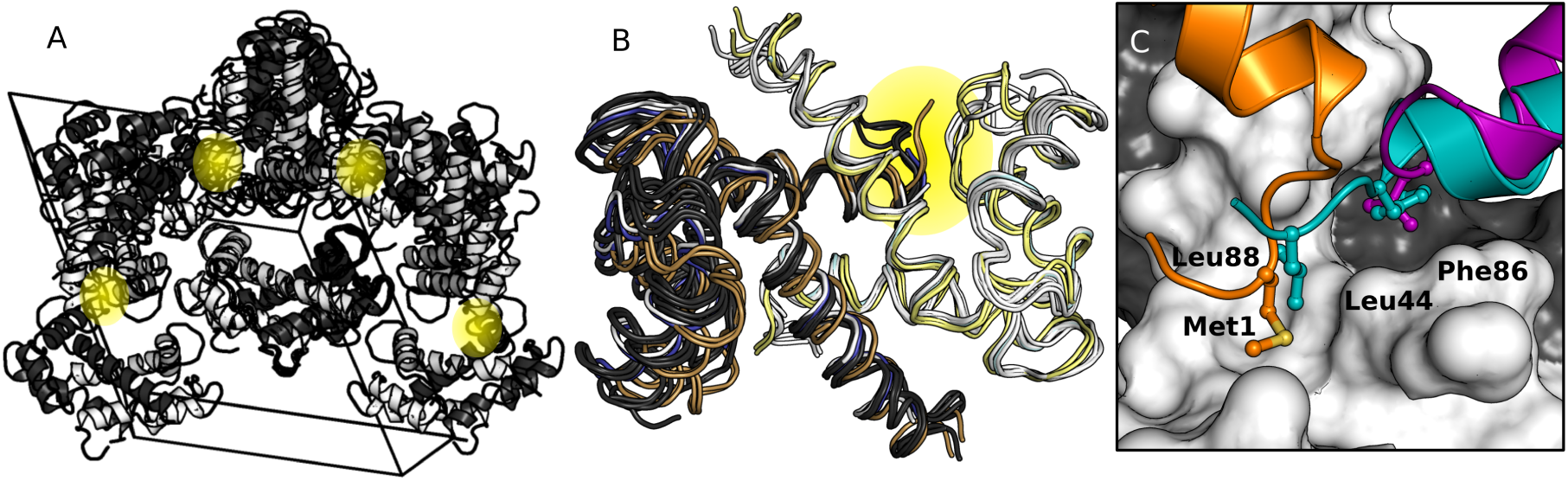
Crystal structure of hA5 reveals variability of peptide interaction surface. A) Unit cell of the hA5 crystal structure showing all symmetry mates. The asymmetric unit consists of 8 homodimers packed together. Peptide binding surface interactions are highlighted with yellow ovals. B) Overlay of all calcium-bound structures of hA5: 1.25 Å crystal structure from this study (white/dark gray, 6 chains), a 2.60 Å crystal structure (PDB: 4dir, cyan/navy, 2 chains), and an NMR solution structure (PDB: 2kay, yellow/brown, 2 chains). Binding site shown in panel C is highlighted in gold. C) The binding site is occupied by crystal symmetry mates in three different configurations (orange, purple, and teal).

Fortuitously, this new structure provides a high-precision view of the peptide binding site participating in three distinct interactions with crystal symmetry mates, thus providing insight into protein-peptide interactions (Fig. 4C). Two largely hydrophobic regions of the binding site coordinate non-specific binding of bulky hydrophobic side chains, with one pocket occupied by either a Met or Leu and the other by Leu or Phe (Fig. 4C). These interactions with crystal symmetry mates likely explain why our attempts at diffusing peptides into the crystal were unsuccessful, as the intra hA5 contacts occlude the canonical peptide binding surface.

The ability of the binding site to accommodate a variety of peptide ligands is demonstrated in that binding is facilitated by occupying either one or both hydrophobic pockets, with few other interactions stabilizing this association. In fact, no intermolecular hydrogen bonds were formed at these interfaces. Across these three binding modes, we also observed the binding site itself changed little (as evidenced by similarity between different chains within the asymmetric unit; Fig. 4B). This perhaps explains the importance of shape complementarity for our binding prediction model.

### Docking models reveal hydrophobic nature of the interaction

Our machine learning model and crystal structure both indicate that binding recognition is mediated by hydrophobic contacts and shape complementarity (Fig 3, Fig 4). To gain structural insight into how this works for individual peptides with radically different sequences, we used ROSETTA to dock peptides to an hA5 dimer extracted from our crystal structure. We docked four peptides known to bind: A5cons (SSFQDWLLSRLP), A6cons (GFDWRWGMEALT), NCX1 (RRLLFYKYVYKR), and *α*-1-syn (GERWQRVLLSLA, see next section). For each peptide, we generated 80,000 docked models by FlexPepDocking,^32^ starting the peptide in several different orientations relative to the protein. We took the top 4,000 models for each peptide and used them for further analyses. To determine the similarity between the resulting models for a given peptide, we clustered them based on their *C*_*α*_ RMSD. Although we allowed up to 50 clusters, 95% of models for each peptide were partitioned into only 5 to 12 clusters.

To summarize the results, we calculated the *C*_*α*_ RMSD between each model and the best-scoring model for that peptide, and then plotted this RSMD against the score for each model (Fig 5A-D). In such plots, the best model appears on the bottom left: the model with the lowest overall score has an RMSD of 0 against itself. We then colored each model on the plot by the cluster it was within.

**Fig 5.**
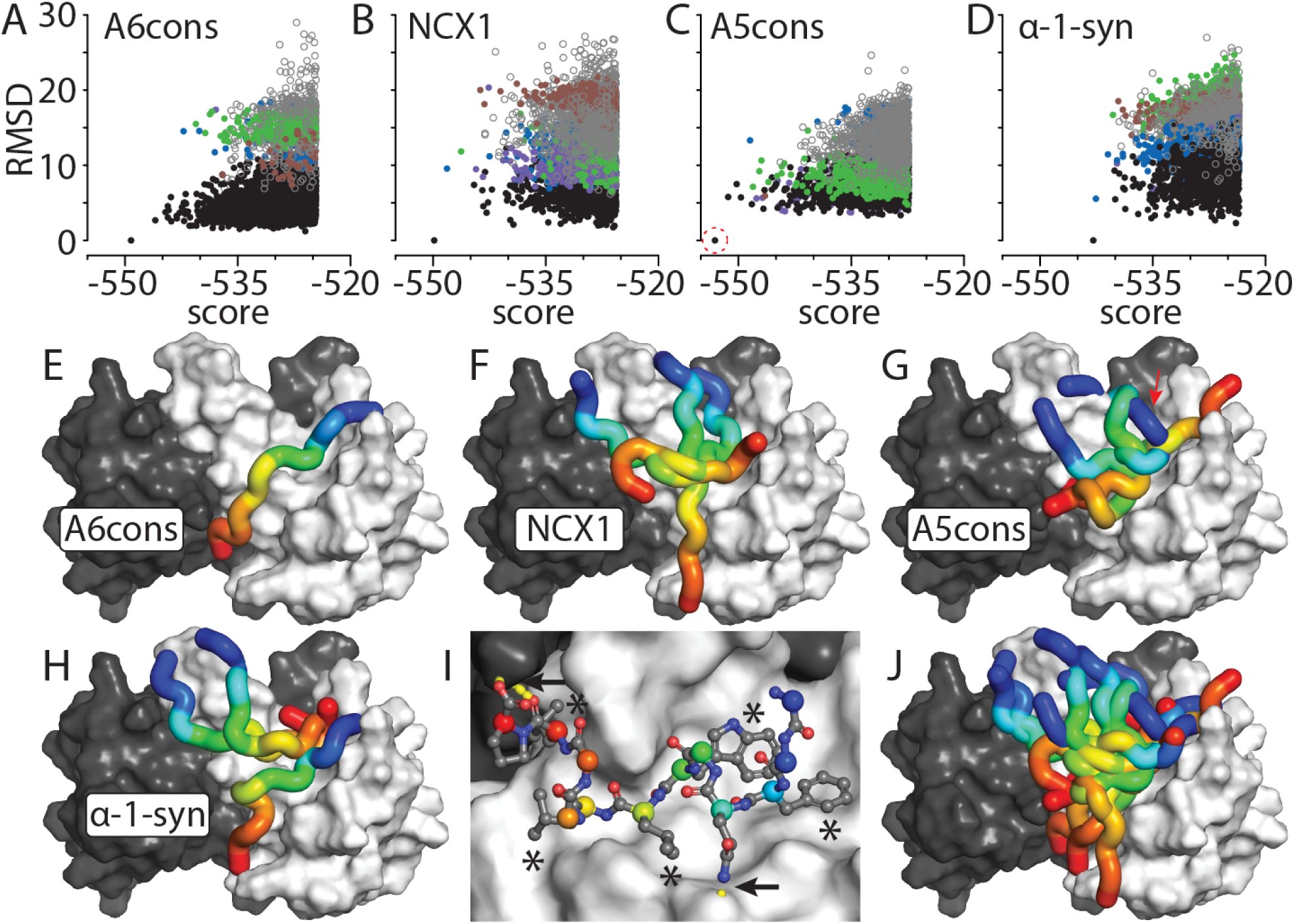
Docked peptides show multiple binding modes. A-D) Docking results for peptides indicated above each graph. Each point is a single model. The color of each point indicates its cluster membership, ranked from the cluster with the best to the worst score: black, blue, green, brown, and purple. Open circles represent peptides taken from clusters besides the top five. The x-axis is the ROSETTA score for the model; the y-axis is the *C*_*α*_ RMSD for each model against the best model for that peptide. E-H) Plausible models for the peptide indicated on the structure. The hA5 input structure is shown as a surface, with chain A and B shown in gray and white. The peptide is shown as a tube, colored from blue (N-terminus) to red (C-terminus). Only the top model is shown for A6cons; the top three models are shown for the remaining peptides. I) Molecular detail of the highest scoring overall peptide model (A5cons). *C*_*α*_ atoms are highlighted with colors matching panel F. The three hydrogen bonds formed between the peptide and hA5 are indicated with arrows; hydrophobic interactions are indicated with “*”. Sidechains that do not interact with S100 have been removed for clarity.J) Overlay of all 10 peptide docks shown in panels E-H.

The A6cons peptide yielded a single, unique, binding solution. Of the top 4,000 models, 70.8% of fell within a single cluster, including the 12 best-scoring models (black points, Fig 5A). For this model, the peptide takes on a largely extended conformation that drapes across the hydrophobic binding surface in a belt-like fashion (Fig 5E). In contrast, the remaining three peptides did not yield unique docking solutions. Take NCX1, for example. The top three models all came from different clusters (the left-most black, blue, and green points in Fig 5B). The peptide in the three models occupies the same basic binding pocket, but it traces through the pocket in three different ways (Fig 5F). The A5cons peptide (Fig 5C, 5G) and *α*-1-syn peptide (Fig 5D, 5H) gave very similar results.

These interactions are almost entirely hydrophobic in nature. As an example, we can look in detail at one of the A5cons models (Fig 5I). This model had the best overall score for any peptide docked to hA5 (red circle, Fig 5C). In this model, the peptide forms five, well-packed hydrophobic interactions (indicated with asterisks), but only three hydrogen bonds to hA5 (indicated with arrows). This dearth of hydrogen bonds is common for all of the peptides. If we average over the cluster containing the best-scoring model for each peptide, A6cons forms the most hydrogen bonds to hA5 (3.2 ± 2), while NCX1 forms the fewest (1.3 ± 1). Thus, as predicted by the machine learning model, polar interactions do not seem to play a key role in defining peptide binding.

We can also use these models to rationalize the finding that many diverse peptides bind. If we overlay the solutions shown in Fig 5E-H onto a single structure, we can see the sheer breadth of structures that are compatible with this binding site (Fig 5J): the interface can accommodate a wide variety of peptide configurations, as long as they can have hydrophobic amino acids and enough flexibility to pack into position.

### The trained model identified a possible new hA5 target peptide

Finally, we attempted to use our trained model to predict new, biologically-plausible targets, for hA5. We used a 12 amino acid sliding window to find 10,477,400 unique 12-mers in 20,206 human proteins extracted from uniprot. Applying our trained model to this k-merized human proteome resulted in a set of predicted interacting peptides (Fig. 6A,B). The resulting distribution of predicted *E* scores is shown in Fig. 6A. The distribution is centered at zero, with a tail extending along the negative (higher enrichment) axis. An estimated 3.9% of proteomic 12-mers had an *E* value below our apparent detection threshold of −1.37.

**Fig 6.**
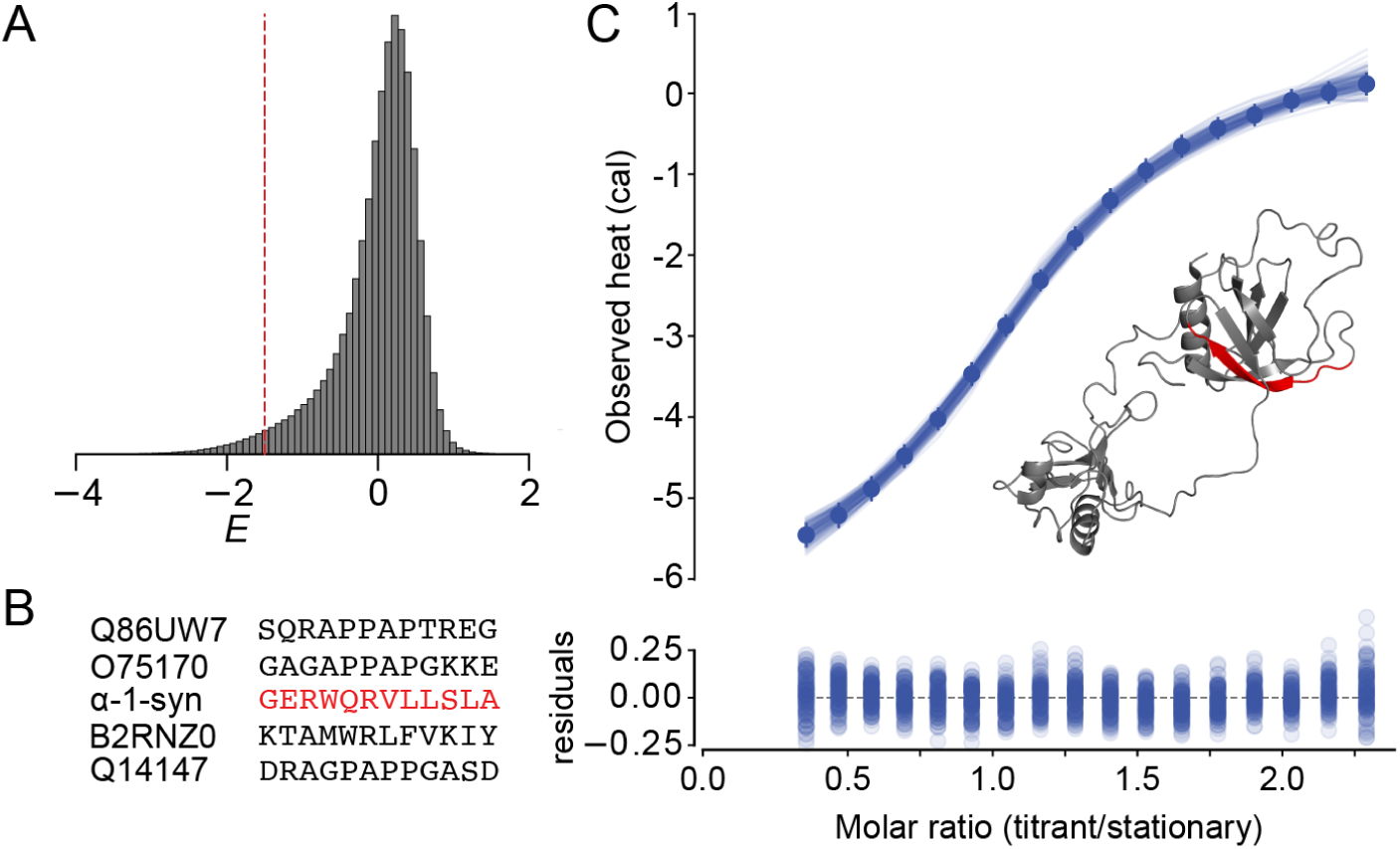
hA5 binds tightly to one of the predicted peptide targets. A) Histogram showing the distribution of E scores for proteomic 12-mers predicted to bind to hA5. Red dashed line indicates the cutoff of *E* = − 1.37. B) Sequences of the five proteomic peptides predicted to bind to hA5. Newly discovered target, *α*-1-syn, is highlighted in red. C) Isothermal Titration Calorimetry (ITC) trace showing binding of peptide *α*-1-syn to hA5. We estimated parameters for a single-site binding model to the data using the Bayesian MCMC sampler in pytc.^33^ Lines show 100 individual fits sampled from the Bayesian posterior probability distribution. Inset shows structure of human *α*-1-syntrophin (PDB entry 1Z87) with the Q13424 peptide fragment (GERWQRVLLSLA) labeled in red. Detailed data on predicted peptides can be found in Table 3.

We next sought to predict specific sequences that would bind. We selected five peptides from the top 0.05% of the *E* score distribution and purchased synthetic versions of these peptides. For experimental tractability, we selected peptides that were predicted to be soluble using the pepcalc.com server. To avoid effects from the peptide termini, we ordered the predicted 12-mer peptides plus 3 additional amino acids taken from the full protein sequence at both the N- and C-terminal ends (Fig. 6A and B). The full peptide sequences and the proteins from which they were taken are shown in Table 3. We measured binding of these peptides to hA5 using ITC. We first conducted all the measurements at 25 °*C*. If we were unable to detect a heat of binding at 25 °*C* we also attempted to measure the interaction at 10 °*C*. Because these protein-peptide interactions are expected to be hydrophobic, we would expect to see non-zero Δ*C*_*p*_ of binding, and thus heats of binding at one or both temperatures.^14,18^

**Table 3:** Model predictions for potential new peptide targets. Model *E* scores and measured binding affinities for peptides predicted by our model to bind to hA5. Flanking amino acids outside the predicted 12-mer are shown in lowercase.

| uniprot accession | Sequence | Protein of origin | $E$ | $K_D(\mu M)$ |
| --- | --- | --- | --- | --- |
| Q86UW7 | agsSQRAPPAPTREGrrd | Calcium-dependent secretion activator 2 | -4.03 | >100 |
| O75170 | dapGAGAPPAPGKKEapp | Serine/threonine-protein phosphatase 6 | -3.94 | >100 |
| Q13424 | gagGERWQRVLLSLAedt | α-1-syntrophin | -4.45 | 5 |
| B2RNZ0 | keiKTAMWRLFVKIYFlqk | Human olfactory receptor 14\|1 | -3.53 | >100 |
| Q14147 | sedDRAGPAPPGASDgvd | TP-dependent RNA helicase DHX34 | -3.88 | >100 |

The peptide extracted from *α*-1-syntrophin protein (referred to hereafter as “*α*-1-syn”) bound to hA5 at 25 °*C* with *K*_*D*_ = 4.8 *µM* (95% confidence, 1.4 to 23 *µM*) and Δ*H* = −4.5 *kcal · mol*^−1^ (95% confidence, −11 to −1.5 *kcal · mol*^−1^) (Fig. 6C). The peptide has little sequence similarity to other previously-identified targets; however, it does possess five hydrophobic residues, including one tryptophan. It also has multiple charged and polar residues that, together, make it readily soluble in water.

The remaining four peptides gave no evidence of binding at either temperature (Table 3). These peptides are quite variable in sequence; however, three of the four (Q86UW7, O75170, Q14147) are rich in proline and alanine and are studded with charged residues. They also notably lack the large bulky tryptophan possessed by the *α*-1-syn peptide (Table 3). Thus, it is possible that these proline rich peptides clash with the binding site despite favorable overall properties. It is less clear what may determine the lack of B2RNZ0 peptide binding to hA5.

## Discussion

We applied a supervised machine learning approach to a previously-measured high-throughput phage display dataset to predict the binding of peptide targets for human S100A5 (hA5). Using this model we were able to: 1) recapitulate the established pattern of specificity for a set of known targets, 2) determine that the major biochemical drivers of peptide binding were hydrophobicity and shape complementarity, and 3) identify a previously unknown target peptide from human *α*-1-syntrophin. By solving a crystal structure of the calcium-bound form of hA5, we were able to propose a biophysical rationale for the low specificity of the protein: there are several different binding modes at the canonical peptide interface. This was confirmed by peptide docking studies, which found that peptides could dock in multiple orientations, while exhibiting a paucity of hydrogen bonds to the hA5 surface. Our results lay the groundwork for a more thorough understanding of the biochemistry and biology of hA5. We also provide evidence that high-throughput binding experiments coupled with deep sequencing and machine learning constitute a potential way forward in understanding the determinants of binding specificity in very low-specificity proteins.

### A step forward in understanding the biochemistry of hA5 specificity

We find that the peptide specificity of hA5 is determined largely by shape complementary and hydrophobic surface area—not polar contacts. These are the most predictive features in our trained model (Fig 3E). This result is further supported by the crystal structure, which shows that interactions between subunits at the peptide-binding surface are mediated by several different hydrophobic contacts (Fig 4C). For example, in one symmetry mate pair, a bulky hydrophobic side chain extends from one symmetry mate into the peptide binding pocket of another. Finally, our docking results show that peptides can be accommodated in multiple orientations in the binding pocket (Fig 5J)—forming many hydrophobic contacts, but few hydrogen bonds (Fig 5I).

As a result, the features that contribute to binding are distributed across the target peptides, rather than being concentrated onto one or two key sites. This observation is a notable deviation from the traditional way of thinking about protein-protein interaction specificity, which is often centered around the idea of binding “hot spots”.^34^ This helps to explain why a straightforward representation of binding preferences as a motif or position-weight-matrix has not been possible for S100 proteins. We suspect that similar patterns may be identified in other low-specificity proteins and that similar approaches to ours may be required to understand the determinants of binding specificity.

While hydrophobicity and shape complementarity are clearly important, our model likely underestimates the contribution of polar contacts. It systematically underestimates the magnitude *E* of both the highest and lowest *E* peptides (Fig 3C). We suspect this is because these values of *E* depend most strongly on specific structural details, rather than the aggregate biochemical features considered by our model. Such contacts may be “smeared out” by the model, and thus make a smaller contribution to the model than they do in the actual molecular interface. This effect must be relatively small, however, as the model performs quite well overall and our structural analyses support a small role for polar contacts at this interface.

### Implications for the biological roles of hA5

The large predicted interaction set for hA5 (Fig 6A) likely reflects the hydrophobic nature of the peptide binding surface. Any peptide that presents a compatible hydrophobic surface is expected to bind, possibly in multiple conformations. Crucially, however, this does not mean that *any* peptide will bind. We found four new peptides that did not interact with hA5 (Fig 6B), in addition to the previously known “SIP” peptide.^18^ Further, we found previously that this specificity has been conserved for hundreds of millions of years in S100A5 paralogs,^18^ suggesting that the low specificity does not represent a total lack of peptide binding preference.

Our results suggest a plausible, but previously unknown target for hA5 (Fig 6C). The peptide we identified is a fragment of human *α*-1-syntrophin, a largely disordered PDZdomain-containing protein that is expressed in a variety of human tissues and serves as a scaffold for various signaling molecules.^35–37^ The peptide fragment is part of a relatively exposed region of the *α*-1-syntrophin PDZ domain, and should be accessible to hA5 in the cell. There are several tissues where both proteins are expressed including kidney and brain.^36–41^ Future biological experiments such as pull-down assays should be used to test whether *α*-1-syntrophin is truly a biological interaction partner of hA5.

Aside from identifying a specific target, our results also allow us to create a rough estimate of the number of putative hA5 peptides that may exist in the proteome. Based on the predictions of our machine-learning model, we estimate that the protein can bind to roughly 4% of the 10,477,400 unique 12-mers in the human proteome. When we sampled five predicted binders, we found that only one bound. If we assume the model yields ≈ 80% false positives when applied to the proteome, there are ≈ 420, 000 potential hA5 targets. If only 10% of these partners are physically accessible—with the rest occluded the interior of proteins or cell membranes—we are still left with 42, 000 peptide fragments that may be expected to bind to hA5.

This suggests that other mechanisms are required to offset the low biochemical specificity of hA5. One possibility is hA5’s precise cellular expression and localization. The protein has a very tight expression pattern and appears to be localized near specific bilayers,^41–43^ thus limiting its available binding targets. hA5 also has relatively low affinities for peptides (≈ *µM*),^18,19^ meaning that both it and/or its partners must be at relatively high concentration for an interaction to form. Finally, it is also possible that there are additional higher-ordered properties of proteins that restrict the true set of possible hA5 target peptides. For example, in addition to the peptide region itself, the nearby regions may need to possess flexibility to accommodate peptide binding—something our peptide model does not take into account.

### Implications for predicting proteomic targets of low-specificity proteins

Finally, our work suggests that even relatively sophisticated machine-learning approaches may not be sufficient to build models that reliably predict new binding targets for low specificity proteins. In our case, only one of the five peptides we sampled from the human proteome interacted with hA5. This low success rate likely arises from a few sources. First, there are errors in the model itself—it does not perfectly reproduce the phage display data. Second, phage-display does not perfectly map to binding of isolated peptides *in vitro*. The third, and likely most important issue, however, is statistical. The total number of non-binding peptides in the proteome is almost certainly very large compared to the number of true targets; therefore, even a small false positive rate in our predictions would cause a huge number of false positives that can swamp out our true predictions.^44^ This means that predicting specific new targets from the proteome, even with an exceedingly accurate model, will be quite challenging. Predictions will thus always require experimental follow up to validate their binding.

### Future directions

Unlike purely sequenced-based methods, our approach provides insight into what biochemical features are recognized by the protein. By recoding the amino acid sequence as a vector of biochemical properties, one gains insight into what features of the amino acids—and the peptide as a whole—are being recognized by the protein, rather than what letters are preferred. This is particularly powerful for a case like hA5, where the amino acid preferences are not obvious, but it will likely be useful for more specific proteins as well. For example, if a motif contains a tryptophan and a tyrosine at a given site, what are the relative contributions of hydrophobicity and hydrogen bonding to binding?

All that is required as input for these calculations is a large collection of peptide sequences with some measured property such as enrichment, binding, or activity. Our software automatically calculates the chemical features and then writes them out in a format that can be fed into any machine learning platform. The approaches we implement here should thus be broadly applicable to other proteins that recognize short linear motifs,^1,2,4^ providing a framework for future studies to decipher the biochemical determinants of binding preferences in these systems.

## Materials and Methods

### Machine Learning Analysis

We implemented our machine learning model in Python 3 extended with numpy,^45^ scipy,^46^ and matplotlib.^47^ We used sklearn 0.21.3 for our random forest regression.^29,48,49^ A full list of the calculated features is shown in Table S1. As noted, some features were calculated using CIDER (using localCIDER 1.7); ^28^ we calculated the remaining features using our own software. We standardized all input features prior to training the model by subtracting the standard deviation and dividing by the mean of that feature as calculated across all observations. We trained the model using the default objective function in sklearn (least-squares). Prior to doing any model fitting, we withheld 10% of the data as a test set. We did k-fold cross-validation on the training data to determine which parameters to include in the fit, using *k* = 10. We determined the relative contribution of each feature to our final trained model using the “feature_importance” method of sklearn, which analyzes node impurity as measured by mean squared error. Our full implementation, including all data files and an example script, is available at https://github.com/harmslab/hops.

### X-ray crystallography

hA5 C43S/C79S was expressed and purified from BL21(DE3) cells as described previously.^18^ To generate crystals, we dialyzed 4 mM protein into 1 mM HEPES, 8 mM *CaCl*_2_, 0.25 mM DTT at pH 7.5. We then mixed this solution 1:1 with 0.2 M (*NH*_4_)_2_*SO*_4_, 20% PEG 8000 (w/v). We grew crystals by hanging-drop at 4 °*C.* We harvested crystals, submerged them in a cryoprotectant solution of 25% PEG 1500, and then flash froze them by plunging into liquid nitrogen.

X-ray diffraction data were collected at the Berkeley Advanced Light Source (ALS) beamline 5.0.3 at cryogenic temperatures on a single high-diffracting human hA5 crystal. Data were processed with iMosflm v. 7.2.1^50^ and scaled with SCALA^51^. Data were cut to 1.25 Å resolution based on the method of Karplus & Diederichs ^52^ with *CC*1/2 > 0.3 and completeness > 50 in the highest resolution bin. Analysis with POINTLESS^51^ indicated space group P3112 as a candidate solution, but molecular replacement trials using PDB structure 4dir and Phaser^53^ failed to correctly solve the structure. Data processing also suggested the data may be twinned, as an L test for twinning gave a score of 0.375^54^. Subsequent molecular replacement trials found a solution in space group P32 with three homodimers (6 chains) in the asymmetric unit. Manual model building was performed with Coot v. 0.8.3^55^ and refinement with Phenix 1.10.1.^56^ In late stages of refinement riding hydrogens were added and TLS was applied with one group per protein chain. The protein chains contain two *Ca*^2+^ atoms each that are well-defined and similar in coordination to previous S100 structures. A few solvent sites showed close approaches and may also be fully or partially occupied metal sites, but in the absence of further evidence these were modeled as waters. Despite the high resolution, the final Rwork/Rfree of the structure was 26.2/29.4 % and the model was unable to be further improved, potentially owing to crystal twinning (Table 2). Nevertheless the binding site interactions are clearly observed in the electron density. The final structure was submitted to the protein data bank as code: 6WN7.

### Docking Studies

Docking analyses were performed using ROSETTA3.1 (build 2018.09.60072),^57^ using the FlexPepDocking binary.^32^ We generated 3-, 5-, and 9-mer fragment libraries using the included ‘make_fragments.pl’ script, with the UNIREF90 database as input. For each peptide, we generated used two starting models, both of which had the peptide in the extended conformation. The models differed in the direction of the chain relative to the binding pocket: *N* → *C* going “up” or “down” the pocket (according to the orientation shown in Fig 5E). When clustered, models came equally from each of the initial docking models, suggesting the results did not depend on the choice of starting model. We executed FlexPepDocking with the flags “-lowres_abinitio -pep_refine -ex1 -ex2aro”. We generated ≈ 80, 000 docked models for each peptide.

After docking, we extracted the top 5% of models (4, 000) for each peptide for downstream analysis. We clustered the models based on peptide *C*_*α*_ RMSD, using hierarchical clustering by unweighted pair group method with arithmetic mean (UPGMA). The cophetic correlation coefficient ranged from 0.7-0.9 for all four peptides. We specified that the software identify 50 clusters; however, we found that 95% of the models ended up in the top 5 to 12 clusters for each peptide. Clustering and data analysis were done Python 3.7 extended by numpy,^45^ scipy,^46^ and matplotlib.^47^ Hydrogen bonds were counted in output structures using VMD, ^58^ with the criterion of < 3.5 *Å*, < 40^°^.

### Isothermal Titration Calorimetry

Synthetic peptides were purchased from GenScript, Inc. For all peptides, we attempted to measure binding at 25 °*C.* If binding could not be detected at 25 °*C* we also attempted the experiment at 10 °*C.* ITC experiments were performed in 25 mM TES, 100mM NaCl, 2 mM *CaCl*_2_, 1mM TCEP, pH 7.4. Samples were equilibrated and degassed by centrifugation at 18, 000 *× g* at the experimental temperature for 35 minutes. Synthetic peptides were dissolved directly into the experimental buffer prior to each experiment. All experiments were performed on a MicroCal ITC-200. Gain settings were determined on a case-by-case basis to ensure quality data. A 750 rpm syringe stir speed was used for all experiments. Spacing between injections ranged from 300s-900s depending on gain settings and relaxation time of the binding process. A single-site binding model was fit to the titration data using the Bayesian MCMC fitter in pytc.^33^ Uniform priors were used for all parameters. The ML estimate was used as a starting guess and the likelihood surface was then explored with 100 walkers, each taking 5,000 steps. The first 10% of steps were discarded as burn in. One clean ITC trace was used to fit the binding model. Negative results were double-checked to ensure accuracy.

## Acknowledgements

We thank Joseph Harman, Anneliese Morrison, and John Muyskens of the Harms lab for their helpful comments on the manuscript.

## Funding

This work was funded by NIH R01GM117140 359 (MJH), NIH 7T32GM007759 (LCW), and NIH F32DK115195-02 (AP). The funders had no role in study design, data collection and analysis, decision to publish, or preparation of the manuscript.

## Availability of data and materials

The code used to conduct the machine-learning analysis is available as a Python package at https://github.com/harmslab/hops. The hA5 crystal structure has been made available on the PDB (ID: 6WN7).

## Author contributions

LCW and MJH conceived the study. MJH secured funding for the work. LCW conducted the ITC experiments. MJH and LCW analyzed the phage display data. MJH developed and applied the *hops* machine learning software. CEW grew the protein crystals. AP conducted the crystallographic data collection, structure solution, and related analyses. LCW, MJH, and AP all contributed to writing the manuscript.

## Competing interests

The authors declare that they have no competing interests.

## Notes

### Competing Interest Statement

The authors have declared no competing interest.

