## Supplementary material for "Learning Peptide Recognition Rules for a Low-Specificity Protein": Table S1

**Table S1: Features used for supervised machine learning.** Features denoted (CIDER) were calculated using the CIDER software package (1). Other features were calculated using our own software package (HOPS: https://github.com/harmslab/hops). Features were organized into the combined feature group categories indicated in the second column.

| feature | Training feature group | Pooled feature group | ref |
|---|---|---|---|
| $\kappa$ | CIDER | polar | (1) |
| $\Delta$ | CIDER | polar | (1) |
| $\Omega$ | CIDER | polar | (1) |
| FER | CIDER | polar | (1) |
| $\Sigma$ | CIDER | polar | (1) |
| dmax | CIDER | polar | (1) |
| $\Delta$max | CIDER | polar | (1) |
| NCPR | CIDER | polar | (1) |
| $F_+$ | CIDER | polar | (1) |
| $F_-$ | CIDER | polar | (1) |
| FCR | CIDER | polar | (1) |
| mean hydropathy | CIDER | hydrophobic | (1) |
| White Interface scale | HOPS | hydrophobic | (2) |
| Engleman scale | HOPS | hydrophobic | (3) |
| % buried in structures | HOPS | hydrophobic | (4) |
| Kyte/Doolittle scale | HOPS | hydrophobic | (5) |
| Octanol scale | HOPS | hydrophobic | (6) |
| Hopp-Woods scale | HOPS | hydrophobic | (7) |
| Uversky scale | HOPS | hydrophobic | (1) |
| cumulative mean hydropathy | HOPS | hydrophobic | (1) |
| side chain accessible area | HOPS | hydrophobic | (8) |
| main chain accessible area | HOPS | hydrophobic | (8) |
| Chou-Fasman, $\beta$ | HOPS | sec. structure | (9) |
| Chou-Fasman, $\alpha$ | HOPS | sec. structure | (9) |
| Chou-Fasman, turn | HOPS | sec. structure | (9) |
| fraction poly-proline II | HOPS | sec. structure | (1) |
| num. hbond acceptors | HOPS | polar | — |
| num. hbond donors | HOPS | polar | — |

| | | | |
|---|---|---|---|
| predicted charge at pH 4 | HOPS | polar | — |
| predicted charge at pH 5 | HOPS | polar | — |
| predicted charge at pH 6 | HOPS | polar | — |
| predicted charge at pH 7 | HOPS | polar | — |
| predicted charge at pH 8 | HOPS | polar | — |
| predicted charge at pH 9 | HOPS | polar | — |
| num. positive amino acids | HOPS | polar | — |
| num. neutral amino acids | HOPS | polar | — |
| num. negative amino acids | HOPS | polar | — |
| net charge | HOPS | polar | — |
| isoelectric point | HOPS | polar | — |
| knob main chain, b | HOPS | geometry | (10) |
| socket main chain, x | HOPS | geometry | (10) |
| socket main chain, y | HOPS | geometry | (10) |
| socket main chain, h | HOPS | geometry | (10) |
| knob side chain, b | HOPS | geometry | (10) |
| socket side chain, x | HOPS | geometry | (10) |
| socket side chain, y | HOPS | geometry | (10) |
| socket side chain, h | HOPS | geometry | (10) |
| side chain volume | HOPS | geometry | (11) |
| molecular weight | HOPS | geometry | — |
| aromatic | HOPS | geometry | — |
